# Minor variations in multicellular life cycles have major effects on adaptation

**DOI:** 10.1101/2022.11.02.514833

**Authors:** Hanna Isaksson, Åke Brännström, Eric Libby

## Abstract

Multicellularity has evolved several independent times over the past hundreds of millions of years and given rise to a wide diversity of complex life. Recent studies have found that large differences in the fundamental structure of early multicellular life cycles can affect fitness and influence multicellular adaptation. Yet, there is an underlying assumption that at some scale or categorization multicellular life cycles are similar in terms of their adaptive potential. Here, we consider this possibility by exploring adaptation in a class of simple multicellular life cycles of filamentous organisms that only differ in one respect, how many daughter filaments are produced. We use mathematical models and evolutionary simulations to show that despite the similarities, qualitatively different mutations fix. In particular, we find that mutations with a tradeoff between cell growth and group survival, i.e. “selfish” or “altruistic” traits, spread differently. Specifically, altruistic mutations more readily spread in life cycles that produce few daughters while in life cycles producing many daughters either type of mutation can spread depending on the environment. Our results show that subtle changes in multicellular life cycles can fundamentally alter adaptation.

**Author summary:** Early forms of multicellular organisms exhibit a wide range of life cycles. Though studies have explored how the structure of a life cycle determines the fitness of early multicellular organisms, far less is known about how it affects their adaptation. Studies that do investigate adaptation typically focus on large scale differences between life cycles, implicitly assuming that at some scale life cycles are similar in terms of their adaptation. In this study we consider this assumption by analyzing adaptation in a class of early multicellular life cycles where the only difference between them is the number of offspring they produce. We use mathematical models and evolutionary simulations to compute the fate of mutations that are either altruistic or selfish, depending on their effects on single cells and the groups to which they belong. We find that despite the similarity between life cycles they can adapt very differently. In particular, life cycles that produce few offspring consistently adapt via altruistic traits, while life cycles that produce many offspring adapt via either altruistic or selfish traits depending on the environment. Ultimately, we find that small scale differences in multicellular life cycles can have large effects on adaptation.

## Introduction

Multicellularity has evolved several independent times over the past hundreds of millions of years and given rise to a wide diversity of complex life [1–11]. A central feature of all transitions to multicellularity is a shift from single cells that reproduce autonomously to groups of cells that can beget group offspring. Although extant forms of multicellularity originated in the distant past, experimental and mathematical models have shed light on possible ways of transitioning from single-celled reproduction to group reproduction [12–14]. In particular, these models have shown that the structure of the multicellular life cycle is instrumental in allowing early multicellularity to fix in unicellular populations [15–17]. Yet these models typically only address the initial appearance of multicellularity when the structure of a multicellular life cycle is the main predictor of success [18, 19]. Far less is understood about how the structure of the resulting multicellular life cycles affect adaptation. We address this subject by comparing the tempo and mode of adaptation in a common class of early multicellular life cycles, in which groups of cells reproduce via fragmentation. Ultimately, we find that minor differences in life cycle implementation can have a significant impact on adaptation and the early evolution of multicellularity.

If we consider the early evolution of multicellularity, there is a diverse array of possible multicellular life cycles [20]. Some switch between distinct unicellular and multicellular phases [21] while others remain strictly multicellular [12]; some include different cell types or species [22, 23] while others are genetically and phenotypically homogeneous [24]; and some are ordered via genetic regulation [19] while others rely on stochasticity [25]. Part of this diversity reflects differences across species, but some of it also emerges from the interplay of chance mutations and selection. For example, the same unicellular lineage can evolve different types of life cycles depending on the timing of selective events— even when exposed to the exact same type of selection [26].

Though we expect that large differences in life cycle structure, e.g. the number of phenotypes, will likely affect adaptation, the impact of more minor differences is less clear. For instance, if two multicellular life cycles are identical in all aspects except the number of offspring they produce, to what extent does that affect the kinds of traits that are likely to evolve? If such minor differences within life cycles are indeed important to adaptation, it may shift the relative weights assigned to the influence of chance versus selection in shaping the evolution of multicellularity.

One distinction among early multicellular life cycles that has been shown to affect adaptation is the mode by which the multicellular group forms or develops [20]. In clonal development the group is formed by cells staying physically attached after reproduction, and in aggregative development the group is formed when cells come together, usually in response to an external stimulus such as a toxin or predator. When groups form through clonal development, cells spend the majority of their time growing and reproducing within groups composed of their kin. As a consequence, kin selection can facilitate the fixation of traits that improve group fitness even if they impose a temporary cost to cells, say through a reduced rate of growth [27–38]. Such altruistic traits are less likely to spread in life cycles with aggregative development, because groups are more genetically diverse and ephemeral which weakens the strength of kin selection. In addition, life cycles with aggregative development often include prolonged periods where cells grow and reproduce independently of any groups, and so any trait that imposes a cost to single cells is less likely to fix [20]. Since altruistic traits may underlie the evolution of cooperation within multicellularity, some have highlighted that their relative inability to fix in aggregative multicellularity may explain why most existing, complex multicellular organisms form groups through clonal development rather than aggregative development [9, 39].

Even when groups form in the same manner other features of the life cycle can affect adaptation. For example, in groups with clonal development the presence of a single cell bottleneck has been shown to promote the evolution of altruistic traits [40–43]. Life cycles with single cell group offspring can use the bottleneck to create phenotypically-homogeneous groups. This allows them to purge cells that exploit other members of the group, which increases the probability that altruistic traits will fix. In contrast, groups without a single cell bottleneck can have difficultly fixing such traits because group offspring will likely be mixes of altruistic and exploitative cells and the altruistic cells are the only ones paying a cost. Indeed the presence of a single cell bottleneck has been identified as a key element in multicellular life cycles that have evolved complex forms of cooperation [44–47]. However, the presence of a single cell bottleneck can have the opposite effect if it occurs at the beginning of a phase of unicellular growth, as occurs in alternating life cycles [20]. In such life cycles altruistic traits are unlikely to fix for exactly the same reason as when groups form by aggregative development, i.e. the cost imposed on cells is actively selected against during the single cell phase. Thus the presence of a single cell bottleneck can either facilitate or inhibit the evolution of altruistic traits, depending on other aspects of the life cycle’s structure.

The contrasting effects of single cell bottlenecks highlights the need for identifying the relevant features of multicellular life cycles that shape adaptation. Rather than focusing on large scale differences in life cycle structure, it may be easier to identify salient adaptive features by adopting a bottom-up approach. To this end, we consider a simple multicellular life cycle in which groups form filaments through clonal development and reproduce via fragmentation. Variation within this life cycle is defined by the single “decision” of how to fragment. We draw inspiration from theoretical models [18, 19] and experimental systems [26, 48, 49] and compare two characteristic ways the fragmentation decision is made in terms of their consequences on adaptation. We find that minor variations— such as the size or number of offspring— can have striking consequences for the types of traits (altruistic or selfish) that fix, the rate at which they fix, and the resulting fitness of the multicellular organism.

## Results

For a model of a simple multicellular life cycle we draw inspiration from filamentous cyanobacteria [10, 50–53]. We consider a life cycle in which cells grow and remain in filaments until reaching a fixed size when they then fragment into smaller filaments. While in principle there are many ways that a group of cells can fragment [18], we study a class of fragmentations in which the group splits evenly, producing daughter groups of the same size. For example, a filament with 16 cells could fragment in four possible ways resulting in different numbers of groups either 2, 4, 8 or 16 (see Fig. 1A). We choose this class of life cycles because it includes well-studied multicellular life cycles [12, 50, 54, 55] such as binary fission and complete dissociation and it also enables a controlled comparison of life cycles where only a single parameter changes (the number/size of daughter groups). We note that this class of life cycle assumes some mechanism [18] or signal [19] that ensures regularity.

**Fig 1.**
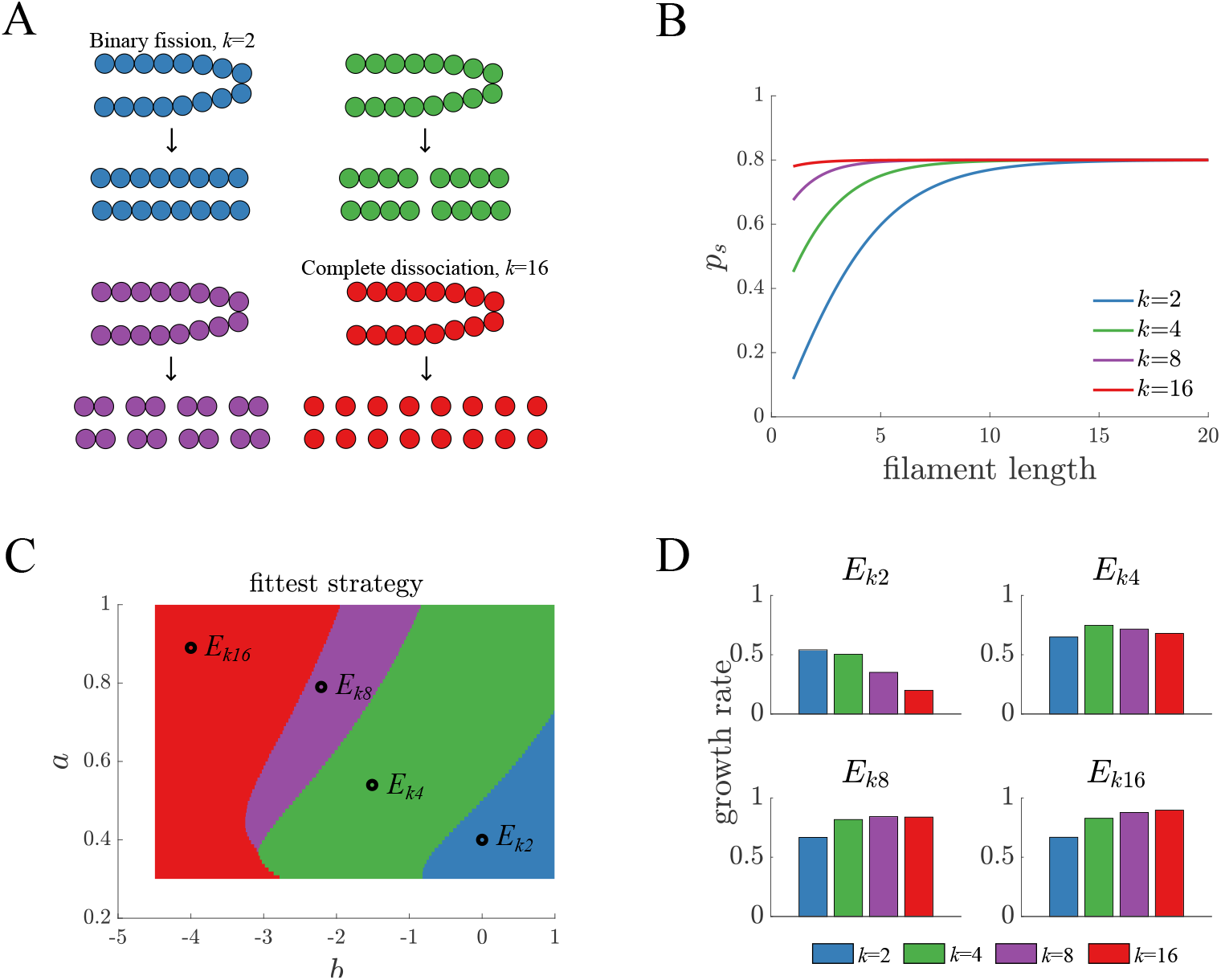
Life cycle fitness across different environments. A) A schematic shows the possible life cycles in which a filament fragments into a set of identical daughter filaments. The parameter *k* indicates the number of daughters and varies between 2 to 16. B) The survival function *p*_*s*_ is shown for different parameter combinations in which each of the life cycles in A) is fittest. For all parameter choices, larger daughters have a higher probability of surviving. C) A fitness landscape shows the *k* that produces the fittest life cycle as a function of the two parameters *a* and *b* in the survival function (see Methods: Life cycles and selection). The color scheme is the same as in panel A and the dots indicate the parameter values used in panel B. D) The long term growth rate for the various life cycles are shown for each of the environments identified in panel C. Binary fission is the least fit life cycle in all environments except *E*_*k*2_.

The fitness of each life cycle is affected by a selection function that imposes a cost to unicellularity. As a result, all multicellular life cycles are fitter than a strictly unicellular life cycle. Selection acts when filaments fragment (similar to [56]) and larger-sized filament offspring have increased survival. The selection function (see Fig. 1B and Methods: Life cycles and selection) imposes different fitness costs on the various multicellular life cycles, and changes in its parameter values can alter which life cycle is fittest. Thus a set of parameter values corresponds to a selective environment that favors a specific life cycle over others. Since the rate of adaptation likely depends on the selective environment, we sought to identify a set of representative selective environments by mapping the fitness landscape for our selection function.

We can determine the fitness for each life cycle by representing it as a branching process and calculating its long-term growth rate (see Methods: Branching process). We find that for a fixed adult size of *N* = 16 cells it is possible to identify environments where each life cycle is fittest (see Fig. 1C, where we use *k* to represent the number of offspring groups). In environments that select for binary fission or complete dissociation, the fitnesses of life cycles are monotonic, e.g. if *k* = 2 is fittest then *k* = 4 is next fittest followed by *k* = 8 (see Fig. 1D). In contrast, in environments where intermediate numbers of offspring are fittest we observed that binary fission was the least fit. This observation led us to focus our analyses on the two extreme values for *k* and their representative selective environments: *k* = 2 for binary fission and its environment *E*_*B*_ (*E*_*k*2_ in Fig. 1C) and *k* = 16 for complete dissociation and its environment *E*_*C*_ (*E*_*k*16_ in Fig. 1C). We note that we performed analyses for the two intermediate environments and intermediate fragmentation patterns (*k* = 4 and *k* = 8) and found they acted similarly to *E*_*C*_ and complete dissociation.

After identifying two types of selective environments, we now consider how each life cycle adapts to these environments. We define a mutant phenotype by the values of two parameters *s*_*c*_ and *s*_*g*_, similar to [20, 57]. The parameter *s*_*c*_ affects the reproductive rate of cells such that when *s*_*c*_ *>* 0 the mutant reproduces faster than its ancestors. The parameter *s*_*g*_ affects the survival of the multicellular filament upon fragmentation such that when *s*_*g*_ *>* 0 the probability of survival increases. Importantly, the interplay between the multicellular life cycle and the values of *s*_*c*_ and *s*_*g*_ determine the fate of the mutant lineage (for a more complete description of how *s*_*c*_ and *s*_*g*_ are implemented in the models, see Methods: Mutant traits).

We first study the fate of a mutant lineage using an analytical approach. If we assume exponentially growing populations then we can express the number of filaments *n*_*g*_(*t*) as a function of time *t* and a characteristic population growth rate *λ*_*g*_,

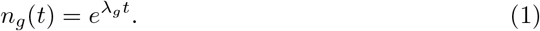

In this expression the value of *λ*_*g*_ determines fitness such that a mutant lineage is fitter than its ancestor if it has a higher value of *λ*_*g*_. Thus, we can solve for the relative fitness *w* of a mutant lineage compared to its ancestor by using a ratio of their *λ*_*g*_ values. If we make a few assumptions concerning how cells and groups reproduce (see Method: Analytical model) we find that

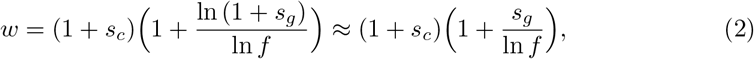

where *f* is the expected number of surviving daughter groups in the ancestor 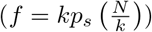, *s*_*c*_ is the mutation’s effect on cell reproductive rate, and *s*_*g*_ is the mutation’s effect on offspring group survival. From this expression we see that the relative contributions of *s*_*c*_ and *s*_*g*_ to fitness depends on the expected number of daughter groups that survive; they are equivalent only if *f* = *e*. Since life cycles that use binary fission can at most result in 2 surviving daughters, the *s*_*g*_ value will always have more of an effect than *s*_*c*_. In contrast, complete dissociation life cycles can span a much greater range in the values of *f*, possibly reaching up to *N*, or 16 in our model. Such high numbers of surviving offspring would make *s*_*c*_ mutations have more of an effect than *s*_*g*_ mutations.

We can use the expression for relative fitness to calculate the rate that a mutant spreads in populations expressing either binary fission or complete dissociation life cycles. We initiate a mutant cell within a filament and compute the growth of populations for a set amount of time, *t* = 20 (see Methods: Adaptation rate). Figures 2 A and B show the resulting mutant proportion of the population when mutations affect either *s*_*c*_ or *s*_*g*_ in isolation. We see that in the binary fission life cycle *s*_*g*_ mutations spread more rapidly than *s*_*c*_ mutations in both *E*_*B*_ and *E*_*C*_ environments. However, in the complete dissociation life cycle we find that either *s*_*c*_ or *s*_*g*_ mutations can spread more rapidly, depending on the environment.

**Fig 2.**
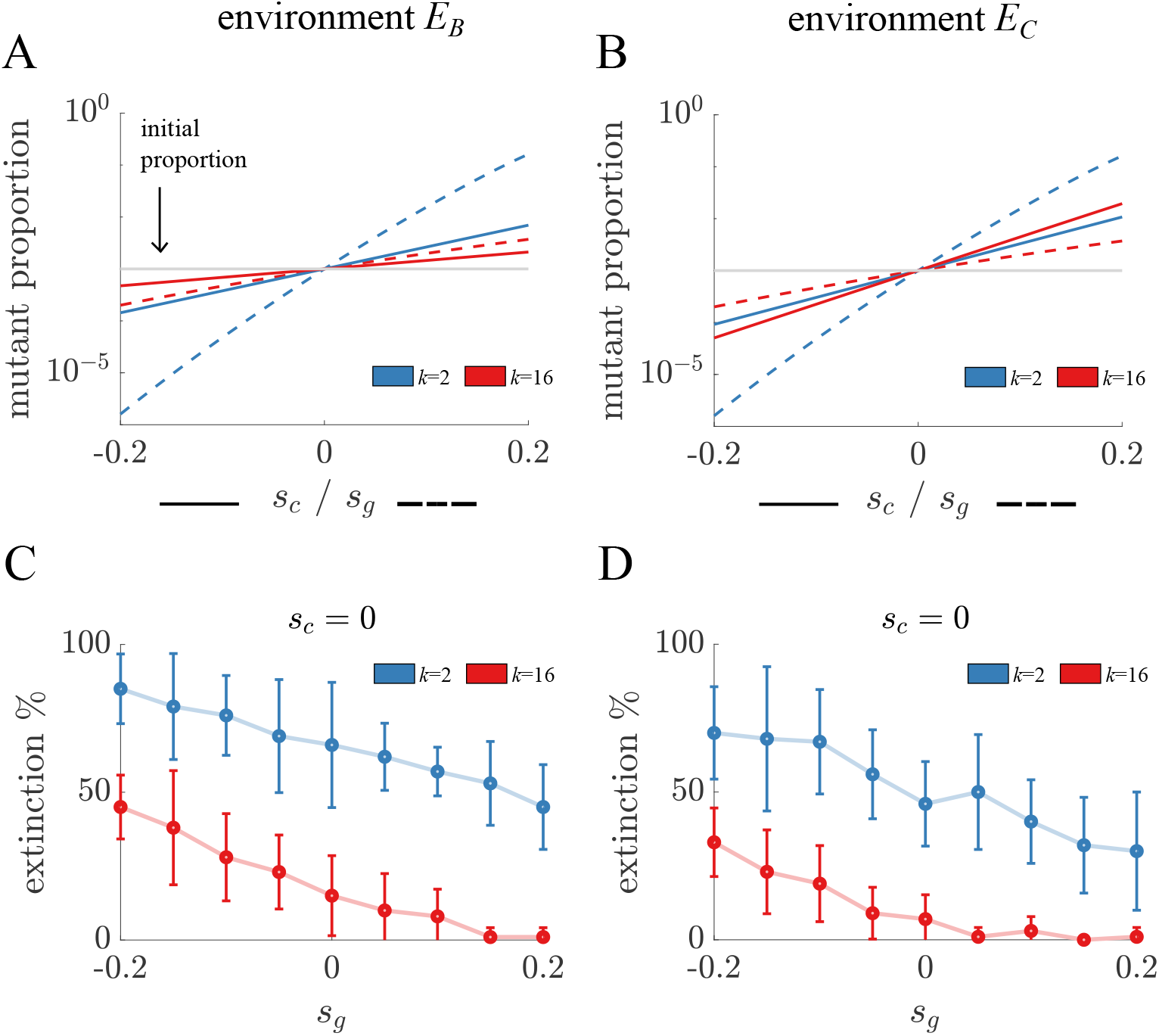
Adaptation and extinction rates for *s*_*c*_ and *s*_*g*_ mutations in isolation. A) The expected mutant proportion at *t* = 20 in *E*_*B*_ is shown for mutations with different values of *s*_*c*_ (solid) and *s*_*g*_ (dashed) in binary fission (blue) and complete dissociation (red) life cycles. The binary fission life cycle adapts faster for both types of mutations, as indicated by the higher mutant proportion when *s*_*c*_ *>* 0 or *s*_*g*_ *>* 0. B) The plot is a companion to A) for the environment *E*_*C*_. The key difference is that *s*_*c*_ mutations spread faster than *s*_*g*_ mutations in the complete dissociation life cycle. They also spread faster than *s*_*c*_ mutations in the binary fission life cycle but not *s*_*g*_ mutations. C) The proportion of mutant lineages that go extinct in *E*_*B*_ is shown as a function of the value of *s*_*g*_ in binary fission (blue) and complete dissociation (red) life cycles. Each point is the mean of 10 samples of 10 simulations and the bars indicate standard deviation. For all values of *s*_*g*_, mutants go extinct more often in the binary fission life cycle. D) The plot is a companion to C) in the *E*_*C*_ environment. Again mutants go extinct more often in the binary fission life cycle—indeed binary fission mutants with the highest value of *s*_*g*_ go extinct more often than all complete dissociation mutants except when *s*_*g*_ *<<* 0.

Another factor affecting adaptation in life cycles is the frequency that mutant lineages go extinct due to random events. Since binary fission life cycles produce fewer group offspring, we may expect that extinctions are more prevalent. We simulate the population growth of a mutant lineage by beginning with a single mutant cell in a filament (see Methods: Stochastic simulations). Figures 2 C and D show the frequency of extinctions in the *E*_*B*_ and *E*_*C*_ environments. In all cases, we observe a higher frequency of extinction in the binary fission life cycle. Even for mutations with the highest value of *s*_*g*_ there are more extinctions in binary fission life cycles then when *s*_*g*_ *<* 0, i.e. deleterious, in complete dissociation life cycles.

After analyzing the spread of *s*_*c*_ and *s*_*g*_ mutations in isolation, we now consider mutations that affect both parameters simultaneously, e.g. epistatic interactions. Although there are many possible interactions between the *s*_*c*_ and *s*_*g*_ parameters, we focus on cases in which there is a tradeoff because they encompass mutations commonly studied in the evolution of multicellularity [57–59]. When tradeoffs exist between cell growth rate and group survival, there are two qualitative types of mutations: 1. selfish mutations, where single cells experience an increased growth rate (*s*_*c*_ *>* 0) and multicellular groups have reduced survival at selection (*s*_*g*_ *<* 0), and 2. altruistic mutations, where single cells experience decreased growth rate (*s*_*c*_ *<* 0) and multicellular groups have increased survival (*s*_*g*_ *>* 0). For each set of *s*_*c*_ and *s*_*g*_ parameters we again calculated adaptation in terms of mutant proportion within each life cycle and assessed extinction rates in each selective environment. Figures 3 - 4 show the results of these calculations for selfish and altruistic mutations, respectively.

**Fig 3.**
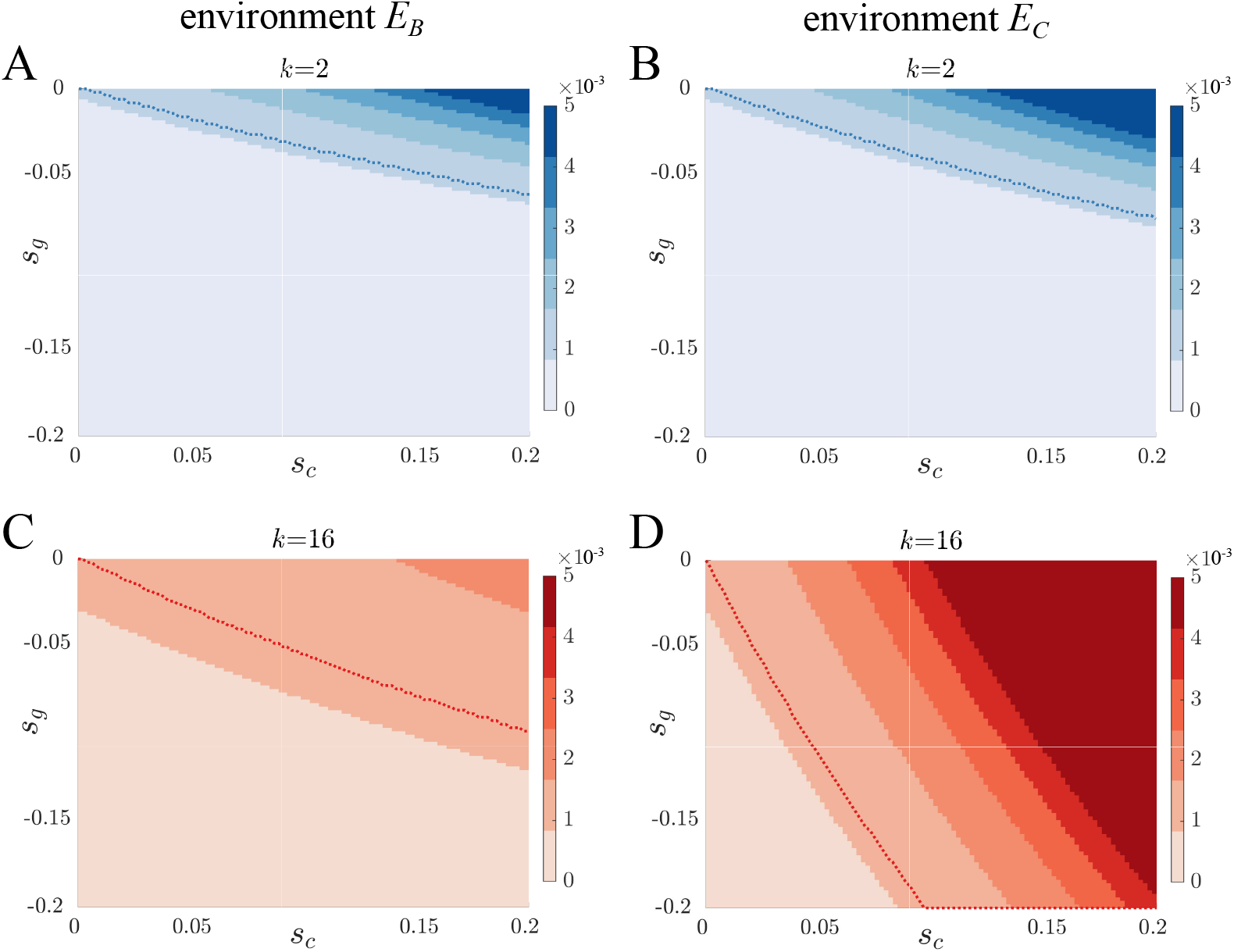
Adaptation via selfish mutations. A-D) Contour plots show the mutant proportion at a fixed time *t* = 20 as a function of the value of *s*_*c*_ and *s*_*g*_ for binary fission (blue) and complete dissociation (red) life cycles in *E*_*B*_ and *E*_*C*_ environments. A greater range of selfish mutations in terms of combinations of *s*_*c*_ and *s*_*g*_ values spread in complete dissociation life cycles. Selfish mutations spread fastest in the complete dissociation life cycle in the *E*_*C*_ environment; however in the *E*_*B*_ environment selfish mutations with a small cost to *s*_*g*_ (e.g. *s*_*c*_ *<* 0.025 for *s*_*g*_ = 0.2) spread faster in the binary fission life cycles. Overall the rate of adaptation for selfish mutations varies more with the environment in complete dissociation life cycles than binary fission.

**Fig 4.**
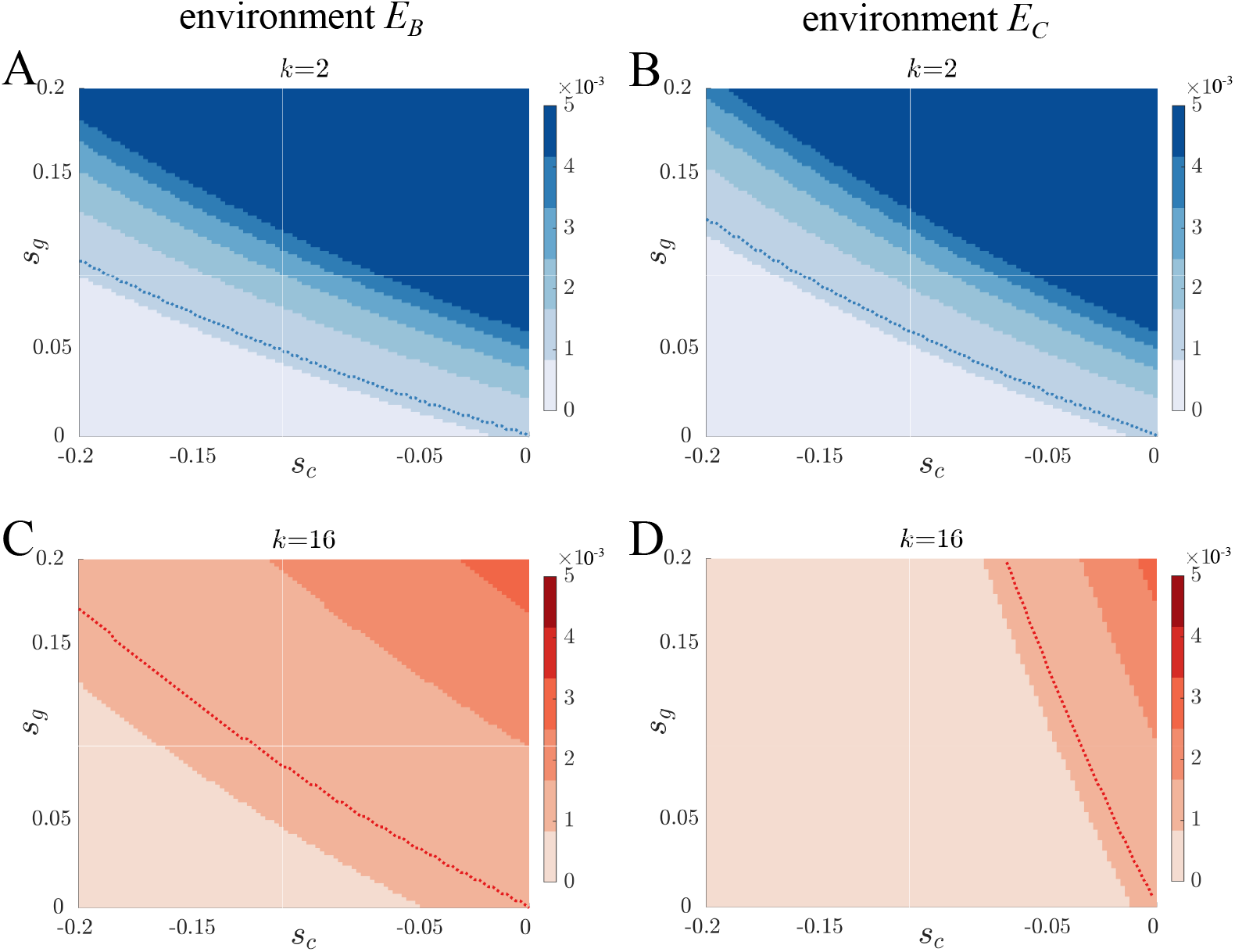
Adaptation via altruistic mutations. A-D) Contour plots show the mutant proportion at carrying capacity as a function of the value of *s*_*c*_ *<* 0 and *s*_*g*_ *>* 0 for binary fission (blue) and complete dissociation (red) life cycles in *E*_*B*_ and *E*_*C*_ environments. Altruistic mutations spread faster and for a greater combination of *s*_*c*_ and *s*_*g*_ values in binary fission life cycles. Again adaptation in complete dissociation life cycles varies more between environments with altruistic mutations spreading faster in the *E*_*B*_ environment.

For selfish mutations Fig. 3 shows that binary fission and complete dissociation life cycles have different patterns of adaptation. Selfish mutations in binary fission life cycles spread similarly in environments *E*_*B*_ and *E*_*C*_. Only selfish mutations with small costs to *s*_*g*_ can increase in proportion. If we compare with complete dissociation life cycles, we find that selfish mutations with higher costs to *s*_*g*_ can spread. Another difference is that the environment has a larger effect on adaptation in complete dissociation life cycles. The area of parameter space in which selfish mutations spread in *E*_*C*_ is more than double that of *E*_*B*_ (77.1% of the area compared to 27.2%).

If we consider altruistic mutations, we also see different patterns of adaptation between the two life cycles (see Fig. 4). The range of parameter space for which altruistic mutations spread is larger in binary fission than complete dissociation in both environments, 76.9% (b.f.) vs 61.3% (c.d.) in *E*_*B*_ and 71.5% vs 18.1% in *E*_*C*_. The rate of spread is also higher such that altruistic mutations can reach proportions in binary fission life cycles that are 31.7 times greater than the highest observed proportion in complete dissociation life cycles. As with selfish mutations, we again observe that adaptation in complete dissociation life cycles is more sensitive to changes in the environment.

In the previous analyses we considered the fate of a single mutation, but it is possible that multiple competing mutations could interact with one another to alter the adaptive process. Thus, we now consider the adaptive consequences of competing mutations by using evolutionary simulations of populations. With multiple mutations competing it takes longer time scales to observe adaptation, and so we simulate populations over repeated rounds of growth and selection similar to serial passaging experiments. In our simulations, populations enact the same multicellular life cycle and grow until they reach a carrying capacity (10^5^ cells), then experience a bottleneck such that 1% of groups survive (see Methods: Stochastic Simulations). When cells within filaments reproduce there is a constant probability (0.01) they will mutate, at which point both *s*_*c*_ and *s*_*g*_ acquire new values from exponential distributions. The resulting evolutionary paths for the mutant trait values from our simulations are shown in Fig. 5 for each life cycle in both selective environments.

**Fig 5.**
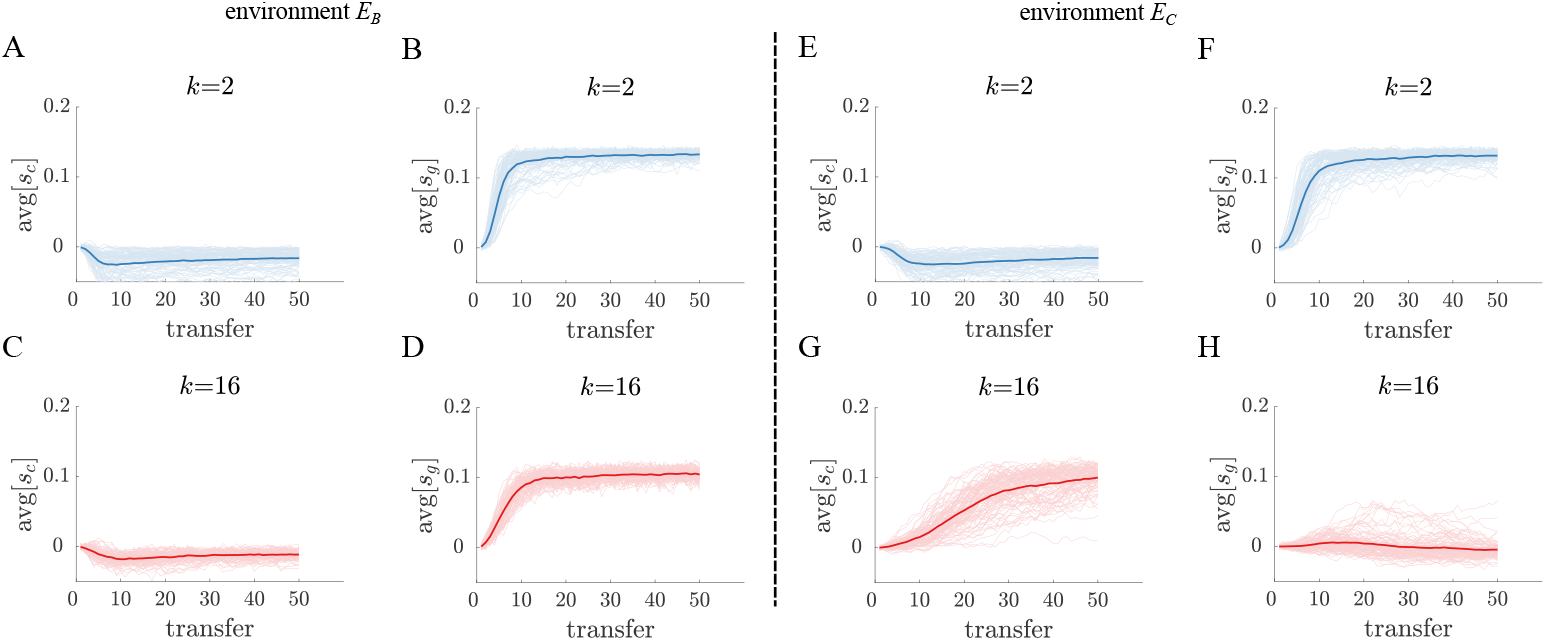
Evolutionary simulations for populations in a serial passage experiment. A-D) Each plot displays the average trait value of either *s*_*c*_ or *s*_*g*_ in 100 independent populations evolving in environment *E*_*B*_. All mutations have a tradeoff such that they are either selfish or altruistic. The thicker lines indicate the average trait value across populations. Both binary fission (blue) and complete dissociation (red) life cycles evolve a higher *s*_*g*_ average at the expense of *s*_*c*_. E-H) These plots share a similar format with panels A)-D) and display trait evolution in the *E*_*C*_ environment.Populations using complete dissociation evolve a selfish trait profile (bottom), while binary fission populations evolve an altruistic trait profile similar to their evolution in *E*_*B*_.

We find that in both selective environments populations evolve to steady state values of *s*_*c*_ and *s*_*g*_—determined by the parameters of our evolutionary simulation —and fluctuate around the steady state thereafter (see Fig. 5). In agreement with our earlier analyses, the majority of populations with binary fission life cycles evolve altruistic traits. In contrast, the complete dissociation life cycle evolved different types of traits depending on the environment. Specifically, complete dissociation evolved selfish traits in *E*_*C*_ and altruistic traits in *E*_*B*_. If we investigate the relationship between *s*_*c*_ and *s*_*g*_ evolved across populations, we find that life cycles can exhibit polymorphic populations (see Fig. 6) in which both altruistic and selfish mutants coexist. Such polymorphic populations are found more often in life cycles with *k* values between binary fission and complete dissociation (see Fig. 6 E). Fig. 6 F shows an example simulation in which selfish and altruistic mutants are stably maintained and regularly swap which makes up the majority of the population.

**Fig 6.**
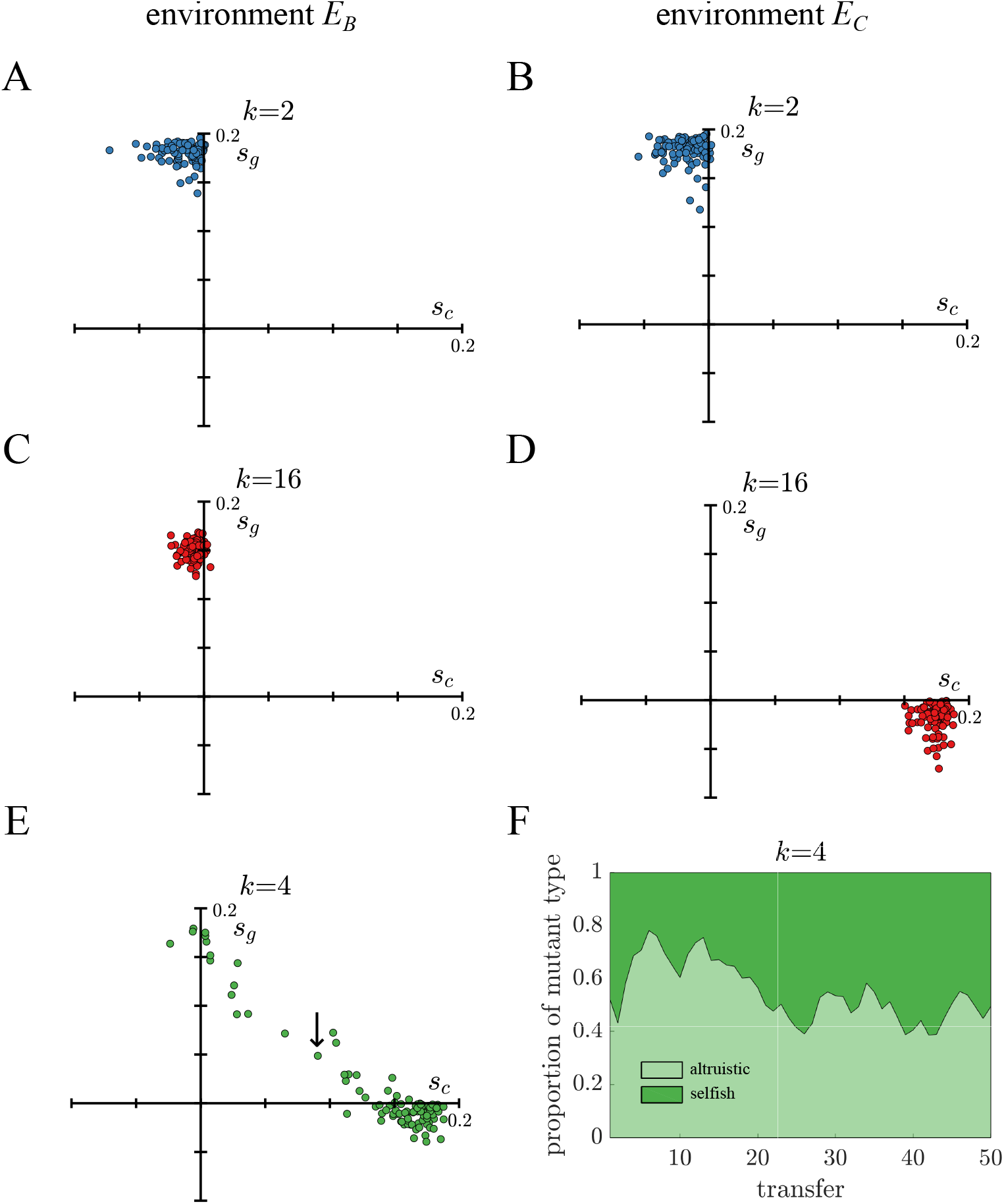
Polymorphisms in populations. A)-D) The average trait values for *s*_*c*_ and *s*_*g*_ are plotted for each independent population in Fig. 5 for binary fission (blue) and complete dissociation (red) life cycles. Populations repeatedly evolved similar altruistic or selfish trait profiles. Although all mutations had a tradeoff with opposite signs for *s*_*c*_ and *s*_*g*_, some populations were polymorphic which allowed both their average *s*_*c*_ and *s*_*g*_ values to be positive. Polymorphic populations were more common for life cycles in their less fit environment, e.g. complete dissociation (red) in *E*_*B*_. E) A similar plot shows the results of evolving a *k* = 4 fragmentation life cycle in *E*_*C*_. Since *k* = 4 is intermediate, lying between binary fission and complete dissociation, its populations are more polymorphic. F) The proportion of selfish/altruistic mutants are shown over 50 transfers for the population indicated by an arrow in E. Altruistic and selfish mutants coexist and regularly swap places as constituting the majority of the population.

Besides qualitative differences in adaptation between the life cycles, there are also quantitative differences in the extent each population adapts. We can define the fitness of a population as the time it takes to reach carrying capacity and thus measure adaptation by the fitness gained in each evolved population. We find that in both environments the binary fission life cycle increased more in fitness than complete dissociation (see Fig. 7), though it was not as fit as the complete dissociation life cycle in *E*_*C*_.

**Fig 7.**
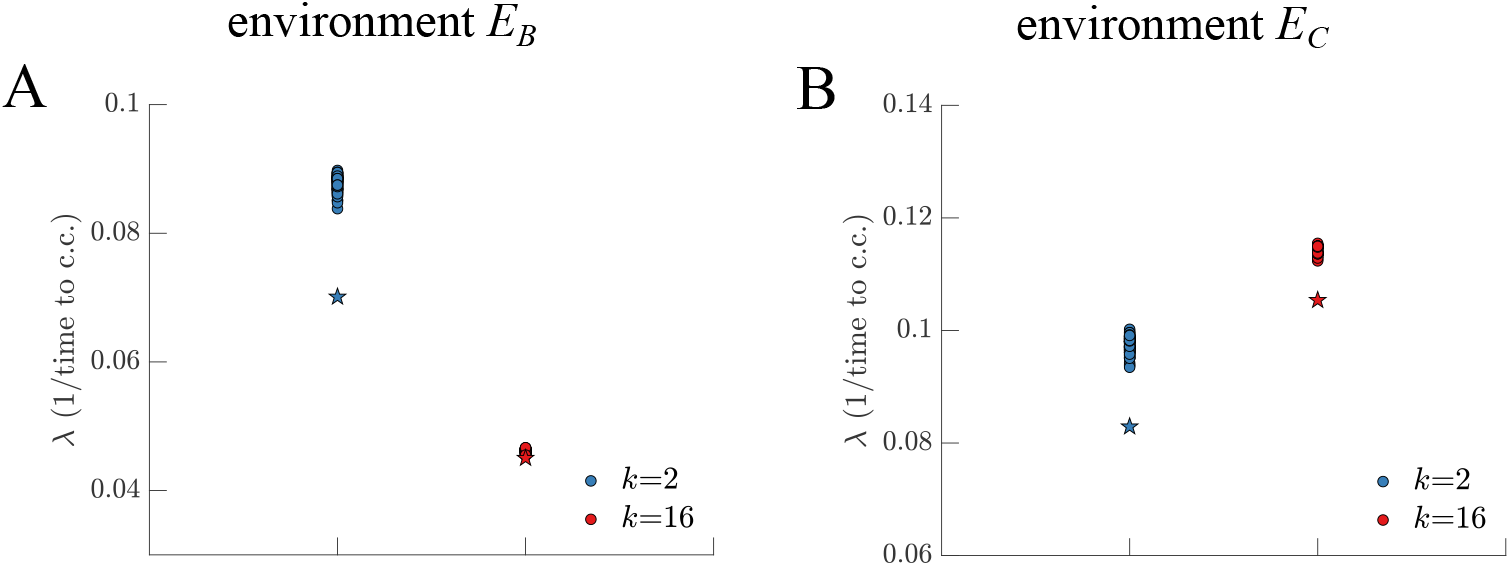
Fitness improvement during evolutionary simulations. A) The scatter plot shows the fitness for evolved populations from Fig. 5 in terms of the time to reach carrying capacity in *E*_*B*_. The star indicates the ancestral fitness prior to evolution. The binary fission (blue) life cycle evolved larger increases in fitness than the complete dissociation life cycle. B) A similar plot to A) shows fitness gains in *E*_*C*_. Again binary fission evolved greater gains in fitness, though it never exceeded any of the complete dissociation life cycle populations.

## Discussion

There are a wide range of ways unicellular organisms can evolve multicellularity. A unifying approach to identify salient differences between types of multicellularity has been to focus on the structure of the multicellular life cycle [13, 16, 17, 20]. Large differences between life cycles have been shown to affect the potential for a nascent multicellular organism to evolve further complexity [15, 20, 39, 44, 57, 60]. Yet, there is an underlying assumption that at some scale, there should be minor differences in a life cycle that do not significantly affect multicellular adaptation. We consider one of the most basic multicellular life cycles, clonal filaments that reproduce via fragmentation, and we use mathematical models to explore the adaptive consequences of subtle variations in fragmentation. We find that qualitatively different types of traits fix across life cycles and at different rates. Our results have implications for the role of historical contingency on the evolutionary trajectories of early forms of multicellularity.

One major finding from our study concerns the role of altruistic versus selfish traits in the evolution of multicellularity. There has been considerable attention devoted to distinguishing forms of multicellularity from social unicellular populations, i.e. multicellular groups from groups of unicellular organisms [61, 62]. A common view is that at first there may be little difference between them but the evolutionary trajectory of a group of cells may lead it to become a more complex form of multicellularity. Features commonly associated with complex multicellularity include those that would not normally evolve in a unicellular context, e.g. altruistic traits that are costly to individual cells but are beneficial to groups [63–66]. In contrast, selfish traits have been associated with a breakdown in multicellularity, e.g. cancers that undermine cell cooperation by reverting to unicellular forms [67–72]. Whether altruistic or selfish traits fix—whether an early multicellular group is evolving away from its unicellular past—is often connected with large-scale differences in multicellularity such as the structure of groups or how they form, e.g. clonally or aggregatively. Instead, we consider the same type of multicellular group with life cycles that differ in only how fragmentation is implemented and find this small change significantly alters whether altruistic or selfish traits fix.

From our models we identify two general factors driving the different evolutionary trajectories of selfish vs altruistic adaptation. The first factor is the expected number of surviving group offspring, which is determined by the survival selection imposed by the environment. When the expected number of surviving group offspring is low *<* ≈ 2.7 improving group survival is more beneficial to fitness than improving cell reproductive rate. Since binary fission life cycles can not yield more than two surviving group offspring, traits improving group survival will always have a relative advantage over traits improving cell reproductive rate. In contrast, complete dissociation life cycles can produce many more group offspring which shifts the balance towards traits that improve cell reproductive rate, even at a cost to group survival.

The second factor driving the difference between altruistic and selfish adaptation is population turnover. When a mutant first arises, its fate depends not only on how it affects life cycle fitness but whether it can survive long enough for those effects to manifest. In binary fission life cycles there is only one population doubling before group reproduction which means that in the vast majority of scenarios there will be only one group offspring with mutant cells. Strong survival selection acting on offspring groups will cause many groups with mutants to go extinct regardless of how the mutations affect cell reproduction. If the mutations improve group survival then this benefit immediately increases the probability the mutant lineage will survive. In contrast, complete dissociation life cycles have log_2_ *N* cell doublings before group reproduction, which provides more opportunity for mutations that affect cell reproductive rate to act. Moreover, the number of mutant cells in a filament prior to reproduction determines the number of mutant offspring groups. Thus, mutations that increase the reproductive rate of cells also increase the probability that the mutant lineage survives, unlike in binary fission life cycles.

We expect that these two factors also play a role in the adaptive differences between broader classes of multicellularity. As an example, we consider the difference between aggregative and clonal multicellularity, in which groups are formed by either coming together or staying together, respectively [73]. A recent experimental study explored the differences in multicellular adaptation between yeast that form groups via aggregation compared to yeast that remain clonal [74]. One finding from this work was that mutant lineages were much more likely to go extinct in the clonal life cycle compared to the aggregative life cycle because of its smaller effective population size. Although the experimental system is quite different from our theoretical models, the aggregative life cycle acts similarly to our clonal life cycle that reproduces via complete dissociation. This similarity likely stems from the fact that aggregative multicellularity typically features a population that assembles randomly to form groups, which allows a nascent mutant population to associate with multiple groups and thereby reduce the risk of extinction. Thus, a salient factor driving the difference in multicellular adaptation between various life cycles may not be how groups are formed but instead the extinction risk faced by any mutant lineage.

Finally, the results of our study indicate that chance and historical contingency can play an important role in the evolution of multicellular complexity [75–77]. In the selective environments we considered, all multicellular life cycles are fitter than a unicellular life cycle. Thus, we expect that if any of the fragmenting life cycles considered here arose in a unicellular population they would outcompete and eventually replace their unicellular ancestors. Yet, following their emergence subsequent multicellular adaptation would likely depend on minor variations within those life cycles and the selective environments. Using simple models we observe that populations would differ not only in whether altruistic or selfish traits took hold but also in the extent that they are polymorphic. Ultimately these results show that chance events in the origins of multicellularity can affect later adaptation and our interpretation of that adaptation.

## Methods and Materials

### Life cycles and selection

We model the population dynamics of filamentous multicellular organisms as simple chains of reproducing cells that fragment when they reach a fixed adult size. We assume that cells reproduce via binary fission and modify their connections accordingly [78]. Once filaments reach adult size, *N* = 16 cells, they split into *k* evenly sized daughters by severing the appropriate connections between cells. We assume that cells do not die in order to sever links— unlike other models [49, 79, 80]. Following fragmentation, all daughter filaments experience selection such that larger daughters have a higher probability of surviving. The function that specifies this size-based selection is *p*_*s*_(*x*) which changes with filament size *x* according to

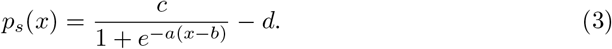

The parameters *c* and *d* were set to 1.7 and 0.9 respectively to ensure *p*_*s*_(*x*) *<* 1 and that there is still room for mutations with *s*_*g*_ *>* 0. The remaining parameters, *a* and *b*, describe the slope and translation of the survival function and vary across different selective environments. Parameter values were selected to guarantee 0 *< p*_*s*_(*x*) *<* 1.

### Branching process

We compute the long-term growth rate of each life cycle by using a continuous time Markov Process. We construct a transition matrix *M* in Eq. 4 where an element *m*_*i,j*_ corresponds to the transition rate from state *i* to state *j*. Specifically, each column tracks the results of a cell division and each row represents the current filament size. The last column displays the results of fragmentation, which happens immediately when the adult size is reached. The size of new daughter filaments is 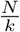 and *p*_*s*_ is the survival function based on offspring size. We can derive the long-term growth rate of a population of filaments by calculating the dominant eigenvalue of the matrix *M*. By varying the parameters *a* and *b* in the selection function *p*_*s*_ (see Eq. 3) we can calculate fitness in different selective environments. For our analysis we pick *a* = 0.4 and *b* = 0 for the *E*_*B*_ environment where binary fission has highest fitness, and *a* = 0.89 and *b* = −4 for the *E*_*C*_ where complete dissociation has highest fitness.

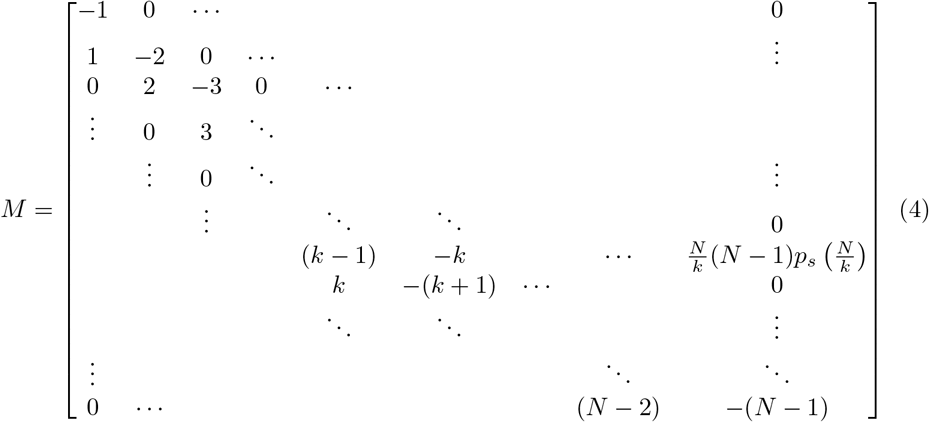

### Mutant traits

In order to study adaptation we let mutations be defined by two trait values: *s*_*c*_ and *s*_*g*_. The *s*_*c*_ trait modifies cell growth rate according to 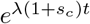 so mutations with *s*_*c*_ *>* 0 cause cells to grow faster than their ancestors. In contrast to *s*_*c*_, which acts on cells outside of their group context, *s*_*g*_ acts on groups as a whole by adjusting their survival. For filaments that are homogeneous and contain mutated cells with *s*_*g*_ *>* 0 the whole filament experiences an increased probability of survival: 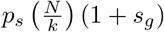. In cases where a filament contains cells with different values of *s*_*g*_ we use the average value for the group’s survival. We constrain values of *s*_*c*_ and *s*_*g*_ within [0.2, 0.2] so that the probability of survival is less than 1.

### Analytical model

We can calculate the population growth of a mutant lineage *n*_*g*_(*t*) as the number of filaments produced per time:

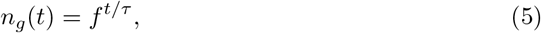

where *f* is the average number of surviving filament offspring produced each iteration of the multicellular life cycle and *τ* is the time it takes to complete the life cycle. We can rewrite this expression in a more conventional form:

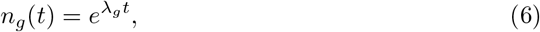

where *λ*_*g*_ is the characteristic population growth rate and is a function of *f* and *τ, λ*_*g*_ = ln (*f*)*/τ*. Since both *f* and *τ* are functions that could depend on many factors, we need to make a few assumptions in order to produce analytical expressions for them. First, we assume that cell populations grow exponentially according to 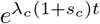 so that the time to complete one multicellular life cycle is *τ* = ln *k/*(*λ*_*c*_(1 + *s*_*c*_)). We note that *λ*_*c*_ and *λ*_*g*_ are not the same term, because *λ*_*g*_ includes the loss of filaments during fragmentation while *λ*_*c*_ does not. Second, we assume that when filaments fragment the probability that an offspring group survives is determined by its size and the value of *s*_*g*_ according to (1 + *s*_*g*_)*p*_*s*_(*x*)—where *x* is the number of cells in offspring groups. We can relate the number of daughters *k* to the size of daughter groups *x* if we assume that all filaments reproduce at some fixed adult size, *N* cells. Putting this together, the number of surviving group offspring produced through a single iteration of the life cycle is 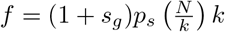 and as a result

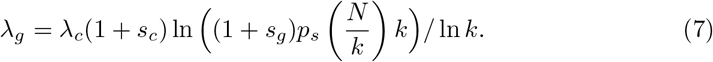

### Adaptation rate

Based on the population growth rate *λ*_*g*_ we can assess adaptation by calculating the mutant proportion for a set of *s*_*c*_ and *s*_*g*_ values. The population starts with an initial mutant proportion *p* and we assume that populations grow with the growth rate as in Eq. 7. We can then define the mutant proportion as

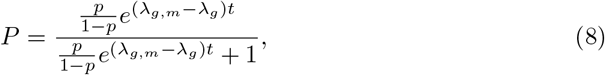

where *λ*_*g,m*_ denotes the growth rate of mutants. From Eq. 8 we see that the factor (*λ*_*g,m*_ − *λ*_*g*_) is the determining factor for adaptation and sets the time scale for evolution. With this factor we can analytically calculate the effect each of the mutant traits have on adaptation. We do this by simplifying the expression (*λ*_*g,m*_ − *λ*_*g*_) to

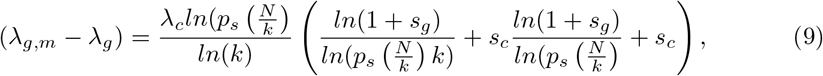

where *λ*_*c*_ is the cell growth rate and 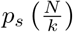 is the selection function that depends on the daughter size 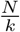. By setting *s*_*g*_ = 0 in Eq. 9 we get the effect from *s*_*c*_ mutations as

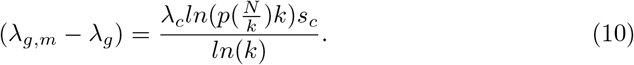

By instead setting *s*_*c*_ = 0 we get the effect from *s*_*g*_ mutations as

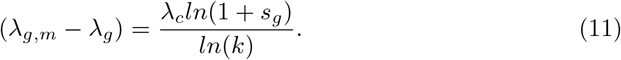

For our computations of the mutant proportion in Fig. 2, 3 and 4 we set *λ*_*c*_ = 1, the growth time to *t* = 20 and the initial mutant proportion to 1*/*1000.

### Stochastic simulations

As a complement to our analytical studies we also performed stochastic simulations tracking the growth of populations of cells. In these simulations cells grow in filaments and fragment according to the fragmentation patterns *k* = 2, 4, 8, 16 when reaching adult size *N*. Selection is applied upon fragmentation and a filament survives if 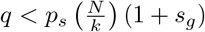, where *q* is a uniform random number. Times for cell division events are sampled from a normal distribution with mean 1 and standard deviation 0.1 to prevent cell reproduction from being completely synchronous. For mutants the reproduction time is scaled by *s*_*c*_ such that the mean is 1*/*(1 + *s*_*c*_).

To estimate the extinction rates we initiate a mutant in a filament of size 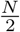. We then simulate the growth of the population and stop once the population reaches 1000 cells. Based on other simulations we found that if the mutant is still alive when there are 1000 cells, it rarely goes extinct afterwards. Indeed the vast majority of extinction events occur within the first iteration through the life cycle.

We structured evolutionary simulations as serial passage experiments. Populations have a set life cycle and grow until reaching a carrying capacity of 10^5^ cells. Upon reaching carrying capacity 1% of the filaments are selected to reseed the population. All simulations begin with a clonal population but at each cell division there is a probability *p*_*m*_ = 0.01 that a mutation occurs in either the mother or daughter cell [59]. If a mutation occurs we draw the value for *s*_*c*_ and *s*_*g*_ from exponential distributions such that *Exp*(*λ* = 20) with equal probability of being positive or negative. We limit the values of *s*_*c*_ and *s*_*g*_ to be within [−0.2, 0.2] so that the probability of survival will not reach 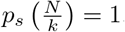

